# *Phanerochaete chrysosporium* strain B-22, a parasitic fungus infecting *Meloidogyne incognita*

**DOI:** 10.1101/622472

**Authors:** Bin Du, Yumei Xu, Hailong Dong, Li Yan, Jianming Wang

## Abstract

We characterized the parasitism by strain B-22 of *Phanerochaete chrysosporium* on the eggs, second stage juveniles (J2), and adult females of the root-knot nematode (*Meloidogyne Incognita*). Strain B-22 had a strong lethal effect against *M. incognita* J2. The highest corrected mortality was 71.9% at 3 × 10^8^ conidia mL^−1^. The estimated LC_50_ value was 0.96 × 10^8^ conidia mL^−1^. Strain B-22 parasitized *M. incognita* eggs 2 days after treatment, causing the depression and dissolution of egg shells. The fungal spores parasitized J2 by gathering in the body wall, germinating to develop hyphae, and crossing the juvenile cuticle to dissolve it, thereby causing the shrinkage and deformation of the juvenile body wall. The spores and hyphae also attacked adult females, causing the shrinkage and dissolution of their bodies and leakage of contents in 5 days. Results of greenhouse experiments showed that different concentrations of *P. chrysosporium* effectively controlled different life stages of *M. incognita* and root knot symptoms in tomato plants. Moreover, the control efficacy increased with increasing conidial concentration; the best results were achieved with 3 × 10^8^ cfu mL^−1^. In the roots, the highest inhibition rate was 84.61% for adult females, 78.91% for juveniles, 84.25% for the egg mass, and 79.48% for the gall index. The highest juvenile inhibition rate was 89.18% in the soil. Meanwhile, strain B-22 improved the plant growth. Thus, *P. chrysosporium* strain B-22 is safe for tomato plants while effectively parasitizing *M. incognita*, making it a promising biocontrol agent against *M. incognita*.

## Introduction

Plant-parasitic nematodes are very common in controlled-environment agriculture and cause economic losses via reduction in greens quality and quantity. Global agricultural losses caused by plant-parasitic nematodes have been estimated at more than $157 billion [1]. Root-knot nematodes (*Meloidogyne* spp.), which comprise 98 species and parasitize almost every species of vascular plants [2], are the most economically important destructive obligate plant-parasitic nematodes. They occur globally, especially in tropical and subtropical agricultural areas, and cause significant yield losses annually (at least $77 billion) to world crops [3]. *Meloidogyne incognita* is one of the most important species in the genus *Meloidogyne*, and it causes dramatic yield losses in many cash crops (such as tomato) in controlled-environment agriculture, which is the main approach to produce vegetables and an important stepping stone to modern agriculture in China. The intensive production, rich soil fertility, suitable soil temperature and moisture, and lack of effective crop rotation in controlled-environment agriculture provide highly favorable conditions for the growth and propagation of M. *incognita*. After 3~5 years of cultivation under controlled conditions with *M. incognita* infection, the crop yield loss is 20–40% or sometimes even up to 60% [4]. *M. incognita* infects the roots of almost all cultivated plants in controlled-environment agriculture, impedes plants’ uptake of water and nutrients due to the formation of giant cells in the roots, and facilitates infection by soil pathogenic microorganisms. Moreover, *M. incognita* is difficult to control because of its wide host range, short generation time, and high reproductive rate [5]. For example, *M. incognita* can infect 1,700 plant species [6]. In China, most greenhouse-grown vegetables are infected with *M. incognita*, causing annual losses of more than $400 million [7]. For these reasons, *M.incognita* has become a prominent problem in controlled-environment agriculture in China.

At present, prevention and control measures against root-knot nematodes include sanitation, crop rotation, the use of organic soil amendments, trap crops, grafting, fertilization, heat-based methods, cultivation of resistant cultivars, transgenic varieties, chemical control, etc. [8,9]. The application of chemical nematicides is the most extensively used and efficient method for the control of *M. incognita*. However, chemical nematicides pose serious threats to the environment and human health and are costly for growers; therefore, the use of chemical nematicides is increasingly being limited or banned [10]. Thus, developing safe, environmentally friendly, and non-toxic alternative methods effective in controlling *M. incognita* are urgently needed. Biocontrol agents provide an alternative strategy for sustainable *M. incognita* management [11,12].

Fungi are an important group of microorganisms that are abundant in soil, and some of these microbes have been characterized for the biocontrol of plant-parasitic nematodes. Nematophagous fungi reduce nematode density by parasitism, predation, or antagonism. Several species of nematophagous fungi have been isolated from around the world. These fungi include *Acremonium strictum*, *Arthrobotrys robusta*, *Catenaria auxiliaris*, *Dactylella oviparasitica*, *Hirsutella rhossiliensis*, *Nematophthora gynophila*, *Paecilomyces lilacinus*, *Pochonia chlamydosporium*, and *Trichoderma harzianum* [13–19]. Many nematode-parasitic fungi have been extensively reported. On the basis of the mechanism of attack on nematodes, nematophagous fungi can be categorized into four major groups: nematode-trapping, endoparasitic, egg-parasitic, and toxin-producing fungi. Nematode-trapping fungi produce trapping devices or specialized structures, which include adhesive networks, adhesive knobs, constricting rings, and adhesive branches, to capture nematodes. Endoparasitic fungi use their adhesive conidia to infect nematodes. These conidia rapidly germinate into hyphae, which can grow, digest, and penetrate the nematode body wall. Egg-parasitic fungi infect nematode eggshells by specialized pegs or appressoria. Simultaneously, these fungi usually produce extracellular hydrolytic enzymes such as proteases and chitinases that play important roles in disintegrating nematode eggshell layers. Toxin-producing fungi produce toxins to paralyze nematodes, and they produce hyphae that can penetrate through and dissolve nematode cuticles [20].

White-rot fungi are used for the removal of toxic pollutants from wastewater [21] and for the improvement of the environment. They can also decompose organic matter such as grass seeds and pathogens from agricultural soils by composting [22]. White-rot fungi commonly inhabit forest litter and fallen trees; they have a strong degradation potential for organic compounds owing to their extracellular oxidative enzymes such as oxidases and peroxidases. These microbes have been demonstrated to efficiently depolymerize, degrade, and mineralize all components of plant cell walls, including cellulose, hemicellulose, and the more recalcitrant lignin [23]. *Pleurotus ostreatus*, a white-rot basidiomycete, can produce toxin droplets to attack and consume nematodes [24,25]. The gene sequences encoding fruiting body lectins of *Pleurotus cornucopiae* are similar with the lectin of a nematode-trapping fungus [26]. *Phanerochaete chrysosporium* is also a white-rot basidiomycete, which can produce diverse extracellular enzymes in the growing hyphal mass [27]. Moreover, the use of *P. chrysosporium* in soil has been shown to be effective in controlling cut chrysanthemum wilt disease by *Fusarium oxysporum* and in improving plant physiological status [28]. The inoculation of *P. chrysosporium* in greenhouse soil has been shown to help overcome the problems of continuous cropping of cucumber by greatly reducing the occurrence of root wilt and root-knot nematode diseases and reducing the relative disease index of root-knot nematodes [29].

The aims of this study were to evaluate the efficacy of *P. chrysosporium* strain B-22 isolated from soil samples from tomato greenhouses in Taigu, Shanxi Province, China, for the biocontrol of the Southern root-knot nematode *M. incognita* under in vitro and field conditions, and to assess the safety of this strain for plant growth. Our results might provide the basis for the development of *P. chrysosporium* as a bio-pesticide for the control of *M. incognita* in greenhouse-grown vegetables. The results will provide new strategies for the theoretical and practical management of *M. incognita*.

## Materials and methods

### Nematode Inoculum Preparation

Infected tomato roots were collected from a tomato under greenhouse field on Taigu (Shanxi province, China) and single egg mass was cultured on tomato as inoculum to establish nematode population for experiment. The species of nematode was identified as M.incognita on the basis of the morphological and morphometrical characters[30]. Egg masses were directly extracted from infected root galls using 1% sodium hypochlorite (NaClO), and separated eggs was gently washed with sterile water to remove the NaClO [31]. Egg masses were kept in distilled water in dark at 25°C for 48h. Then the hatched juveniles were counted for survival under stereoscopic microscope (Nikon Instruments Inc., Tokyo, Japan). The suspensions of *M. incognita* were diluted to approximately 200 juveniles per milliliter with distilled water. After surface sterilized, the eggs, juveniles and females were stored at 4 °C for subsequent trials.

### Phanerochaete Chrysosporium Inoculum Preparation

*P. chrysosporium* B-22 was obtained from Plant pathology department, Shanxi Agriculture University in China and was cultured with potato-dextrose agar medium. The P. chrysosporium was morphologically identified as Burdsall [32]. Five days after incubation at 25 °C, the purified *P. chrysosporium* B-22 was used to produce spore suspensions for inoculation, then spore suspensions were adjusted to obtain 3×10^8^ cfu mL^−1^ for treatments with autoclaved distilled water and counted using hemocytometer [33].

### Parasitic effect of P. chrysosporium strain B-22 on the eggs, J2, and adult females of M. incognita

Root samples infected with *M. incognita* were collected from greenhouse-grown tomato plants in Taigu, Shanxi Province, China. The root samples were gently washed with sterile water and further processed in the Laboratory of Nematology at Shanxi Agricultural University, Taigu. *M. incognita* egg masses and adult females were directly collected from the root samples. Adult females having surface sterilising with 1% NaCOl were suspended in sterile water and the density adjusted to 100 females mL^−1^. Egg masses were added into sterile petri dishes with sterile water and incubated at 25 °C ± 1 °C for 3 days in the dark to hatch the *M. incognita* J2, which having surface sterilising with 1% NaCOl were then incubated to obtain a 1,000 J2 mL^−1^ suspension. Egg masses were centrifuged in 1% NaOCl at 2,000 rpm for 3 min, and free eggs were collected to prepare a 3,000 eggs mL^−1^ suspension.

The parasitic effects of strain B-22 on the eggs of *M. incognita* were determined as described by Zhang et al. [34]. In brief, 100 µL of *M. incognita* egg suspension (about 300 eggs) was placed in a sterile petri dish containing strain B-22 spore suspension at 3 × 10^8^ cfu mL^−1^. A blank control was prepared with an equal volume of sterile water instead of the strain B-22 spore suspension. Petri dishes were incubated at 25 °C ± 1 °C for 3 days. Samples were taken for microscopic examination using a Nikon 80i microscope with an image analysis system (Nikon Instruments Inc.).

The parasitic effects of strain B-22 on *M. incognita* J2 were determined according to Schwartz [35]. Briefly, 100 μL of *M. incognita* J2 suspension (about 100 J2) was placed in a sterile petri dish containing strain B-22 spore suspension at 3 × 10^8^ cfu mL^−1^; these petri dishes were incubated at 25 °C ± 1 °C for 1 day, after which *M. incognita* J2 were picked up and placed on a prepared glass slide with 4% agarose for examination by fluorescence microscopy with an image analysis system (Nikon 80i microscope). Microscopic examinations were performed 2 days later.

The parasitic effects of strain B-22 on *M. incognita* adult females were determined according to Dong et al. [36]. Briefly, strain B-22 was grown on water agar (WA) until the colony diameter reached near the edge of the dish. Sterile coverslips were placed in the WA, and adult females (about 10 females) were transferred onto the coverslips and incubated at 25 °C ± 1 °C for 5 days. A control group was prepared without strain B-22. Samples were subjected to microscopic examination using the Nikon 80i microscope with an image analysis system.

### Lethal effect of P. chrysosporium strain B-22 on the second-stage juveniles of M. incognita

Second-stage juveniles (J2) of *M. incognita* (about 100 J2 100 µL^−1^) were placed in sterile petri dishes containing 1.875 × 10^7^, 3.75 × 10^7^, 7.5 × 10^7^, 1.5 × 10^8^, and 3 × 10^8^ cfu mL^−1^ of a conidial suspension of *P. chrysosporium* strain B-22, as determined by a Neubauer hemocytometer. Control contained sterile water and a suspension of *M. incognita* J2. Petri dishes were incubated at 25 °C ± 1 °C for 72 h, and samples were taken at 24 h intervals to count the number of dead J2 using a Nikon stereomicroscope (Nikon Instruments Inc., Tokyo, Japan).

The mortality of *M. incognita* J2 was determined by the visual inspection of stiff individuals that stayed motionless after a gentle probe with a bamboo needle. The results were used to calculate the mortality and corrected mortality rates (%) of *M. incognita* J2. The median lethal dose (LC_50_) of strain B-22 conidia for *M. incognita* J2 was determined by simple linear regression analysis of the conidial concentration of strain B-22 and corrected mortality rate among J2 of *M. incognita*: y = ax + b, where “y” is the corrected mortality rate among J2 of *M. incognita* and “x” is the conidial concentration of strain B-22. Mortality (%) = (number of dead J2 in each treatment/total number of test J2) × 100%. Corrected mortality rates (%) = (number of dead J2 in each treatment − number of dead J2 in the control treatment)/(1− number of dead J2 in the control treatment) × 100%.

### Greenhouse evaluation of the control efficacy of *P*. *chrysosporium* strain B-22

*P. chrysosporium* strain B-22 was incubated in potato dextrose agar plates at 25 °C ± 1 °C for 10 days, after which 5 mL of sterile water was pipetted onto the surface of the plates, and the spores were scraped off from the plates [37]. The spore suspension was then filtered through a fine-mesh screen (diameter 0.15 mm) to separate the spores from the hyphae. The concentration of the spore suspension was determined by a Neubauer hemocytometer and then adjusted to 1.875 × 10^7^, 3.75 × 10^7^, 7.5 × 10^7^, 1.5 × 10^8^, and 3 × 10^8^ cfu mL^−1^ and stored at 4 °C until use.

A greenhouse experiment was laid in a completely randomized design with six replications (pots) using autoclaved soil sterilized at 121 °C for 2 h. Soil was individually transferred into plastic pots (4kg soil pot^−1^), and three seeds of the *M. incognita*-susceptible F1-hybrid tomato cultivar JG9002 were planted in each pot. When tomato seedlings were growing, each pot was treated with the above spore suspension at the following concentrations: 1.875 × 10^7^, 3.75 × 10^7^, 7.5 × 10^7^, 1.5 × 10^8^, and 3 × 10^8^ cfu mL^−1^. After 15 days, every pot was artificially inoculated with 1,000 *M. incognita* J2. All pots were maintained in a greenhouse at 25 °C, 16 h sunlight, and 65% relative humidity. Pots were fertilized weekly with about 3 g L^−1^ of Poly Fertisol (N:P:K = 14:10:14) and watered daily as needed. Pots inoculated with *M. incognita* J2 but not with the *P. chrysosporium* strain B-22 spore suspension served as controls, while pots that were not inoculated with *M. incognita* J2 or the *P. chrysosporium* strain B-22 spore suspension served as blanks.

After 10 weeks, plant and soil samples from all treatments were randomly collected and brought to the laboratory for counting the nematodes in the roots and the soil for the calculation of the root knot index. The count of nematodes in the soil was determined as per Castillo et al. [38]. Briefly, nematodes were extracted from 100 cm^3^ samples of soil by centrifugal flotation. Sterilized water was used to wash the soil; then, the filtered water, which was passed through a 710 μm mesh sieve, was collected in a beaker in which it was mixed with 4% kaolin (v/v). This mixture was centrifuged at 1,100 *×g* for 4 min, after which the supernatants were discarded, while pellets were resuspended in 250 mL MgSO_4_, and the new suspensions were centrifuged at 1,100 *×g* for 3 min. Supernatants were sieved through a 5 μm mesh, and nematodes collected on the sieve were washed with sterilized water, transferred to petri dishes, and counted under a stereomicroscope. The nematode count in the roots was determined as in Sharon et al. [39]. Egg masses were dissected from the roots and treated with 0.5% NaOCl. Free eggs were collected and examined. Egg samples (about 100 eggs) were incubated in 1 mL of water for 2 days, and the hatched J2 were counted. The extent of root galling damage was determined [40,41] in treatment and control pots after 10 weeks. According to root gall damage, the severity of tomato root galling was assessed on a scale from 0 to 10, where 0 = no knots on roots and 10 = all roots severely knotted or no root system. Root knot index = Σ(number of diseased plants in each rating × score/total number of plants investigated × highest rating) × 100%. Six replications of each treatment were included.

### Safety of P. chrysosporium strain B-22 on tomato plant growth

Plant height, root length, aboveground fresh mass, and root fresh mass were measured independently in each treatment and control after 10 weeks of growth. Plant height and root length were measured with a graduated ruler, while plant fresh mass was weighed on an electronic balance.

### Statistical analysis

All experiments were repeated six times. Data were processed using Microsoft Excel 2007 and expressed as means ± standard deviation (n = 6). The significance of differences in the counts of females, egg masses, and juveniles in the roots; tomato root gall index; and measurements of plant height, root length, aboveground fresh mass, and root fresh mass were examined using the *t*-test and one-way analysis of variance using SPSS 17.0 (SPSS Inc., Chicago, IL, USA). Statistical significance was considered at p-values of less than 0.05.

## Results

### *Lethal effect of strain B-22 on* M. incognita *J2*

Strain B-22 conidial concentration and treatment timing significantly influenced the mortality of *M. incognita* J2 (Table 1). Compared with the control, the mortality of J2 in these experiments was significantly (p < 0.05) greater in all treatments the strain B-22 suspension at the different conidial concentrations, from 1.875 × 10^7^ to 3 × 10^8^ cfu mL^−1^. Mortality increased linearly with the conidial concentration of strain B-22 and time of treatment. The result was similar for all three treatments (24 h, 48 h, and 72 h). In general, the corrected mortality rate of *M. incognita* J2 24 h after treatment was 42.8%. Moreover, corrected mortality rates of *M. incognita* J2 with the conidial concentration of 1.875 × 10^7^ cfu mL^−1^ was 21.6%. The highest corrected mortality (71.9%) was observed 72 h after treatment with the conidial concentration of 3 × 10^8^ cfu mL^−1^. LC_50_ values decreased with time of treatment (Table 1). Estimated LC_50_ values were 3.6, 2.4, and 0.96 × 10^8^ cfu mL^−1^ after 24, 48, and 72 h of incubation, respectively.

**Table 1.** Effects of treatment timing and conidial concentration of the *Phanerochaete chrysosporium* strain B-22 suspension on the mortality of second stage juveniles (J2) of *Meloidogyne incognita*. Values within columns followed by different lowercase letters are significantly different at 0.05 level.

| Time (h) | Conidial concentration ($\times 10^7$ conidia mL<sup>-1</sup>) | Mortality (%) | Corrected mortality rates (%) | Linear regression equation | Correlation coefficient | LC<sub>50</sub> ($\times 10^8$ conidia mL<sup>-1</sup>) |
| --- | --- | --- | --- | --- | --- | --- |
| 24 | 30 | 50.1 ± 1.0d | 42.8 | Y = 0.7511 × X<br>+ 22.91 | 0.8539 | 3.6 |
| | 15 | 46.7 ± 1.3f | 38.9 | | | |
| | 7.5 | 40.2 ± 1.0g | 31.5 | | | |
| | 3.75 | 33.2 ± 1.4h | 23.4 | | | |
| | 1.875 | 31.6 ± 0.5h | 21.6 | | | |
| | Control | 12.7 ± 1.179i | -- | | | |
| 48 | 30 | 62.0 ± 1.17bc | 54.5 | Y = 0.9246 × X<br>+ 28.19 | 0.9681 | 2.4 |
| | 15 | 53.6 ± 1.2de | 44.4 | | | |
| | 7.5 | 47.6 ± 0.83ef | 37.3 | | | |
| | 3.75 | 41.7 ± 0.65g | 30.2 | | | |
| | 1.875 | 40.1 ± 0.95g | 28.3 | | | |
| | Control | 16.4 ± 0.4i | -- | | | |
| 72 | 30 | 76.6 ± 1.1a | 71.9 | Y = 1.16 × X<br>+ 38.85 | 0.944 | 0.96 |
| | 15 | 65.4 ± 2.4b | 58.4 | | | |
| | 7.5 | 60.2 ± 0.92c | 52.2 | | | |
| | 3.75 | 50.6 ± 0.9def | 40.6 | | | |
| | 1.875 | 48.9 ± 0.6f | 38.6 | | | |
| | Control | 16.7 ± 0.35i | -- | | | |

### Parasitic effect of strain B-22 on *M. incognita* at different life stages

*P. chrysosporium* strain B-22 quickly parasitized the eggs of *M. incognita* after 2 days of treatment. In the initial period of parasitism, the spores of strain B-22 were in contact and conglutinated with the eggs, and they geminated and produced short hyphae around them (Fig. 1a). Strain B-22 grew rapidly, causing the aggregation of the inner contents of the eggs. The hyphae of strain B-22 ran through the eggs, causing shrinking of the egg shell. More hyphae grew from the egg (Fig. 1b). Over the following 2 days, the egg shell was deformed and shrunk further. This continued until the egg shell was completely dissolved by strain B-22. The eggs were broken by the hyphae. At last, the dense mycelia of strain B-22 enveloped the egg, which at the time looked abnormal and misshapen (Fig. 1c). In the controls that were not inoculated with the conidia of strain B-22, the eggs of *M. incognita* were intact; microscopic examination confirmed that they had a smooth surface and uniform contents (Fig. 1d).

**Fig. 1.**
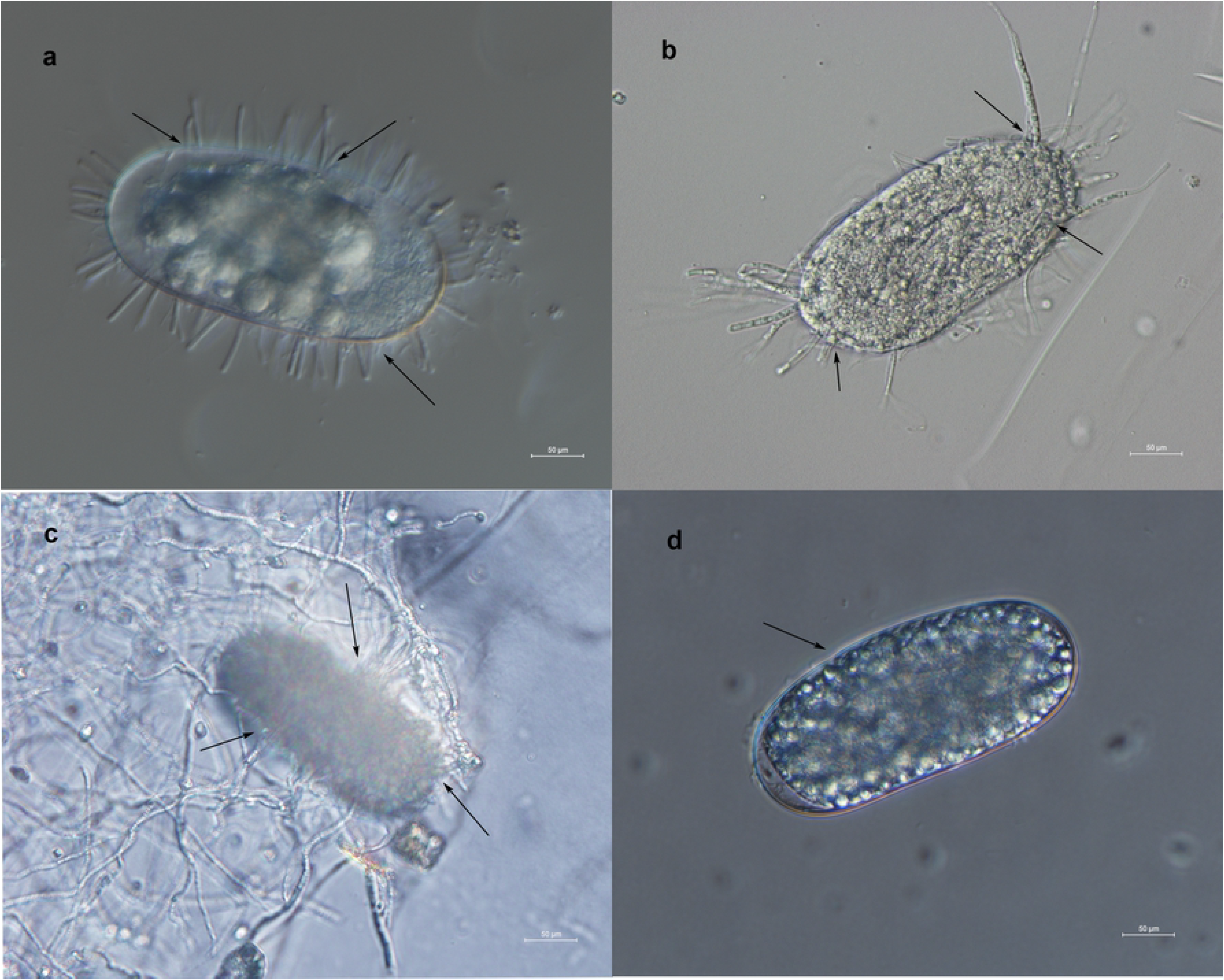
Parasitism by *Phanerochaete chrysosporium* strain B-22 on the eggs of *Meloidogyne incognita.* (a) Spores germinated, and short hyphae developed around the egg; (b) hyphae penetrated across the egg shell; (c) the egg shell was dissolved. (e) Control: healthy, intact eggs. All observations were under 40× magnification.

Strain B-22 also parasitized *M. incognita* J2. After 3 days of incubation, the spores of strain B-22 were seen on the cuticle of J2 upon microscopic examination, as observed in the case of parasitism on eggs. Strain B-22 conidia gathered in the body wall of the J2 (Fig. 2a). Spores geminated and produced hyphae from the body of J2. Thus, strain B-22 grew on the surface of the cuticle of J2 of *M. incognita* (Fig. 2b). With time, strain B-22 produced more mycelium on J2 and dissolved their cuticles, causing shrinkage and deformation of their body wall (Fig. 2c). After 5 days of incubation, microscopic examination showed that the J2 cuticles were dissolved or severely deformed. Dissolved residues of the bodies of J2 were seen clearly. The J2 cuticle line was bent and shrunken (Fig. 2d). Eventually, strain B-22 produced massive spores to develop a dense mycelium, and parasitized the body surface of the J2. The color of the cuticle was extensively altered (Fig. 2e). In untreated controls that were not inoculated with the conidia of strain B-22, the *M. incognita* J2 showed an intact body wall and slow movement (Fig. 2f).

**Fig. 2.**
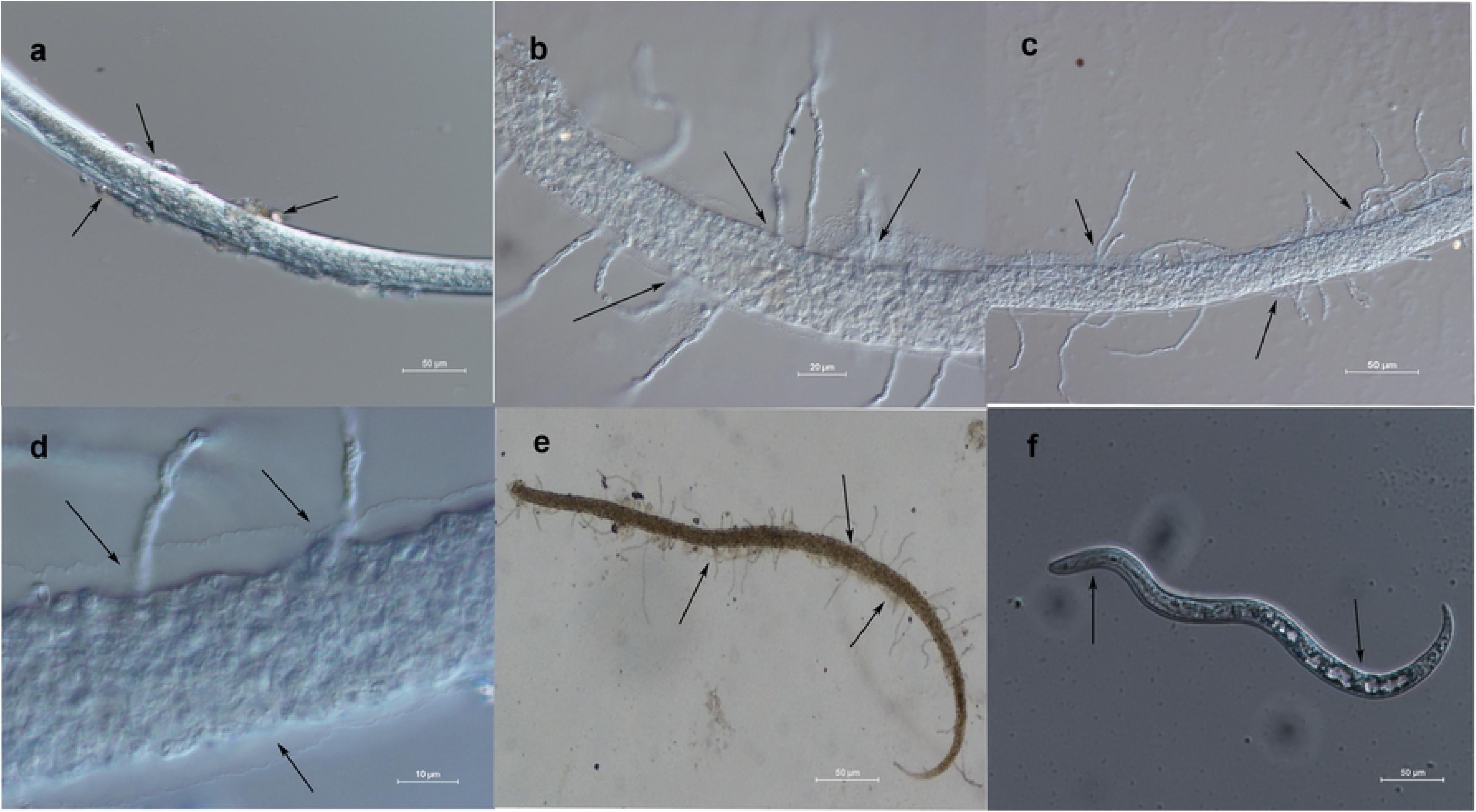
Parasitism of *Phanerochaete chrysosporium* strain B-22 on the second stage juveniles (J2) of *Meloidogyne incognita.* (a) Spores were in contact with J2 (20× magnification); (b) spores geminated, and hyphae grew out from the bodies of J2 (40× magnification); (c) more mycelium was produced from the bodies of J2 (20× magnification); (d) the cuticle of J2 was bent and shrunken (40× magnification); (e) J2 were parasitized by strain B-22 hyphae (20× magnification). (f) Control: healthy J2 (20× magnification).

Strain B-22 also parasitized *M. incognita* adult females. Upon 4 days of incubation, the spores of strain B-22 surrounded the adult females and stuck to their surface. The contents of the bodies of the females leaked out, and short hyphae grew out from their bodies (Fig. 3a). After 7 days of treatment, strain B-22 formed dense hyphae crossing the body walls of adult females. Moreover, the adult female body looked severely atrophied and was partly dissolved due to leakage (Fig. 3b). In the untreated controls that were not inoculated with the conidia of strain B-22, *M. incognita* adult females showed complete and healthy bodies with a smooth surface and obvious boundary. Inside, the contents of the adult female body were intact (Fig. 3c).

**Fig. 3.**
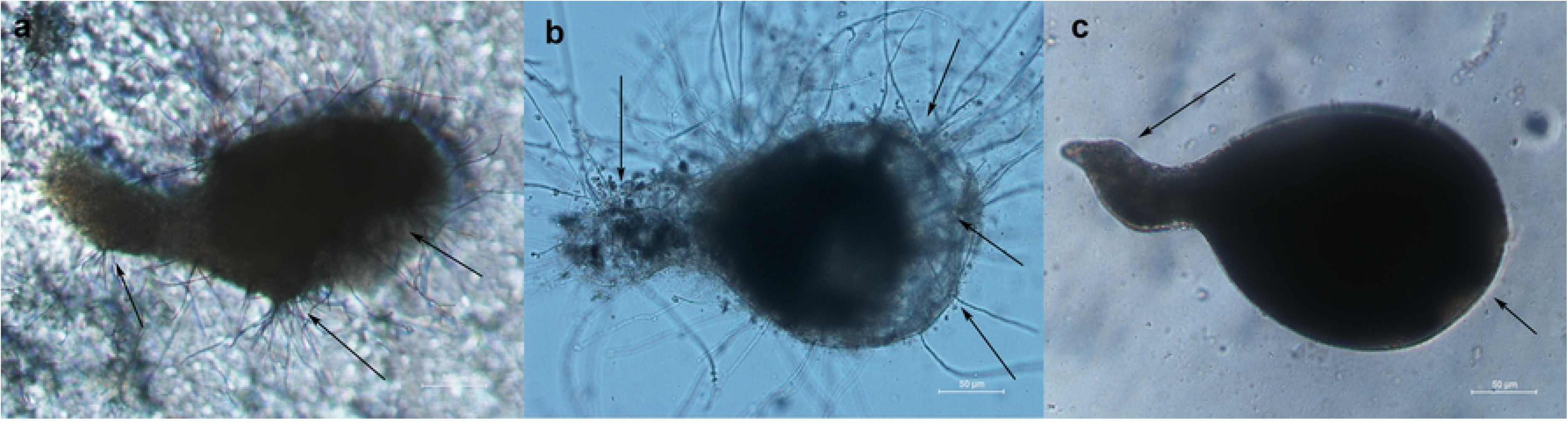
Parasitism of *Phanerochaete chrysosporium* strain B-22 on adult females of *Meloidogyne incognita.* (a) Spores geminated, and the contents of the adult female bodies leaked out; (b) dense hyphae crossed the adult female body, which was partly dissolved. (c) Control: healthy adult female. All observations were under 10× magnification.

### Greenhouse evaluation of the control efficacy against root galling by strain B-22

The conidial concentration of strain B-22 significantly influenced the different stages of *M*. *incognita* (Table 2). The antagonistic effect of different conidial concentrations on the nematodes was significantly (p < 0.05) greater in the treatments than in the blanks and controls. *P. chrysosporium* strain B-22 significantly decreased the quantities of J2, females, and egg masses of *M. incognita*, as well as the root gall index. The number of J2 in the soil also decreased due to treatment with strain B-22. In general, the control efficiency of *M. incognita* significantly increased with increasing conidial concentration of *P. chrysosporium* strain B-22. The inhibition ratio of the females ranged between 46.15% and 84.61% in the roots, while that of the juveniles ranged from 45.57% to 78.91%, and that of the egg mass was from 50.39% to 84.25%. Furthermore, the inhibition ratio of the root gall index was 33.33–79.48%; and in the soil, the inhibition ratio of the juveniles reached 68.51–89.18% (Table 2).

**Table 2.**
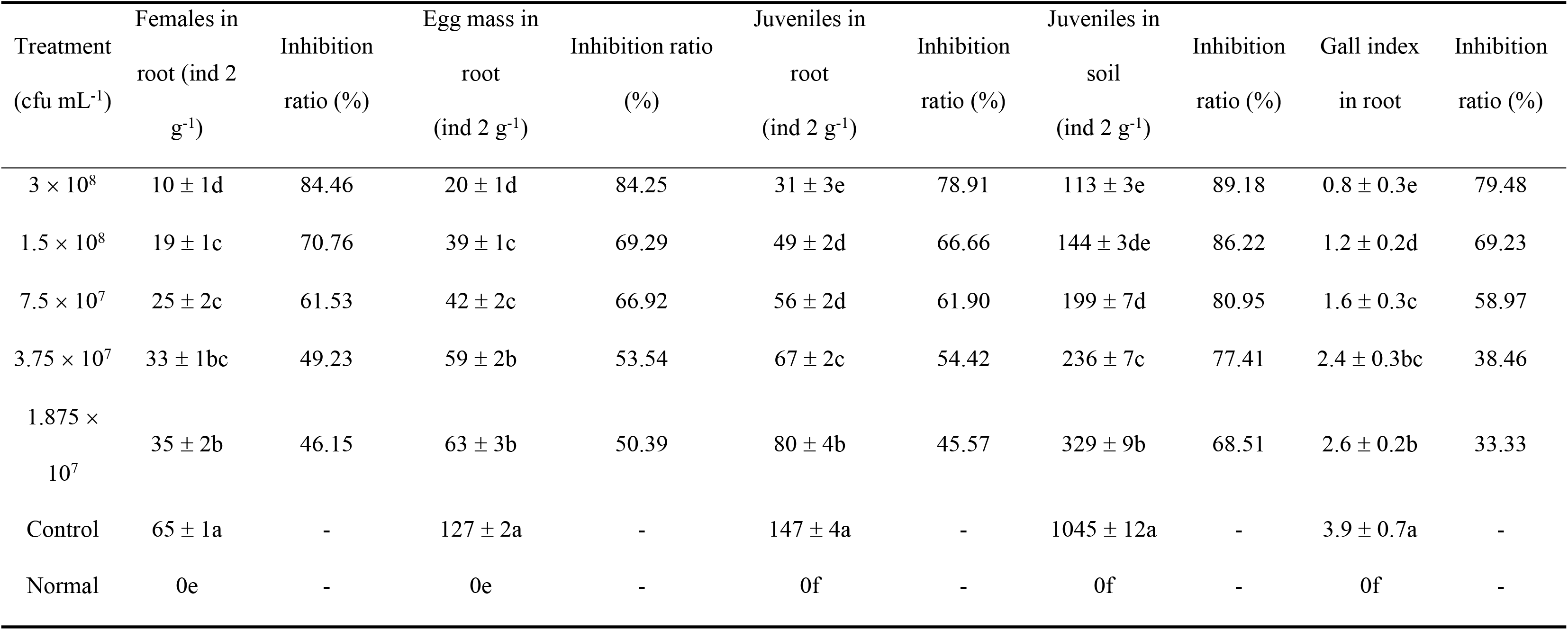
Controlling effects of *Phanerochaete chrysosporium* strain B-22 at different concentrations against *Meloidogyne incognita.*Values within columns followed by different lowercase letters are significantly different at 0.05 level.

### Safety of strain B-22 for tomato plant growth

The use of *P. chrysosporium* strain B-22 in soil was found to be safe for tomato plant growth, and in fact, it had a growth-promoting effect. The conidial concentration of strain B-22 significantly influenced the promotion of plant growth in tomato (Table 3). Plant height, root length, aboveground fresh mass, and root fresh mass were significantly (p < 0.05) greater upon treatment with *P. chrysosporium* at different conidial concentrations compared with the blanks and controls. The growth-promoting effect on tomato plants increased with increasing conidial concentration. The highest promotion was achieved by a conidial concentration of 3 × 10^8^ cfu mL^−1^. In this case, plant height was 29.5 cm; root length was 30 cm; aboveground fresh mass was 9 g; and root fresh mass was 1.5 g, while the corresponding values in the controls were 9.7 cm, 9.1 cm, 1.5 g, and 0.28 g, respectively. The highest increase rates for plant height, root length, aboveground fresh mass, and root fresh mass were 202.06%, 185.71%, 426.66%, and 292.85%, respectively (Table 3).

**Table 3.**
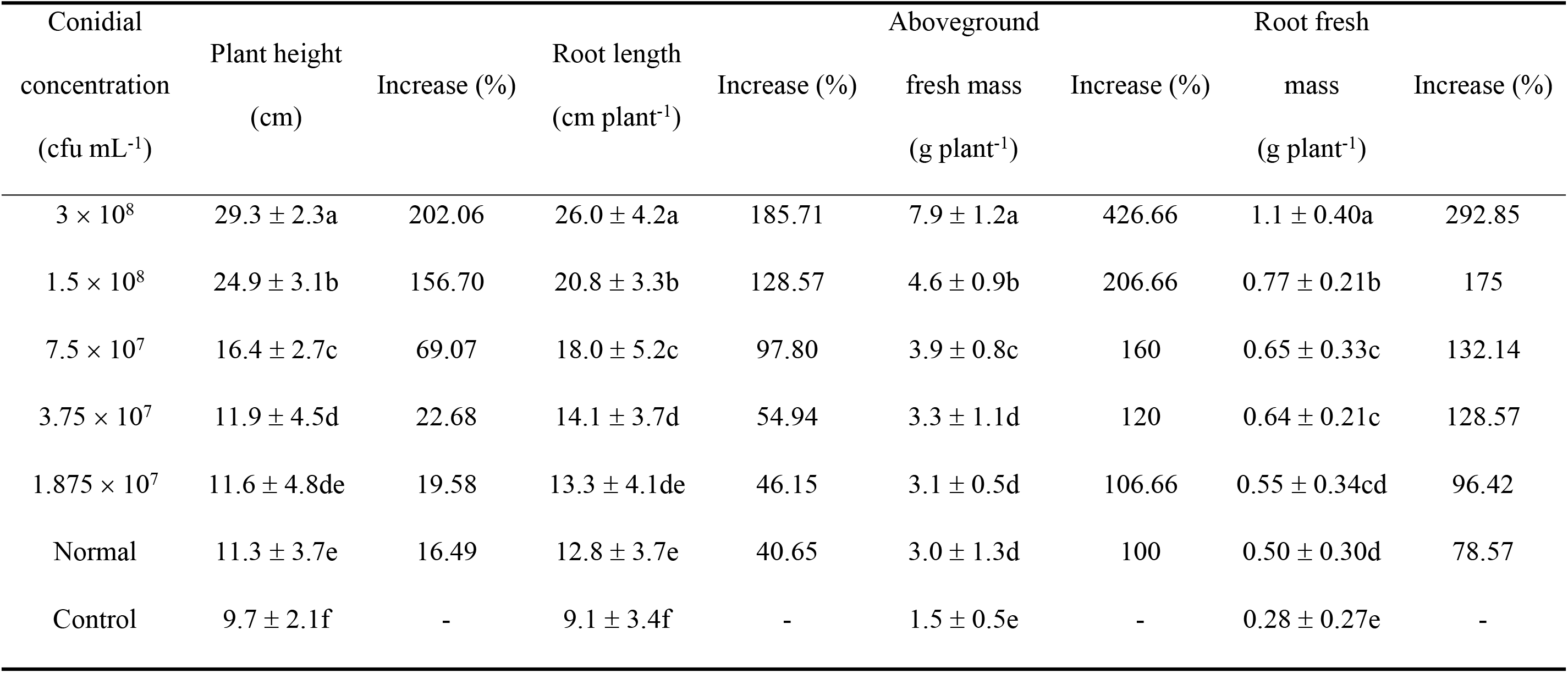
Effects of *Phanerochaete chrysosporium* strain B-22 at different concentrations on tomato plant growth. Values within columns followed by different lowercase letters are significantly different at 0.05 level.

## Discussion

*P. chrysosporium* is a white rot fungus. Some other such fungal genera include *Trametes*, *Bjerkandera*, *Pleurotus*, *Fomes*, *Polyporus*, *Poria*, and *Coriolus*. *Pleurotus ostreatus* has been identified as a nematophagous fungus [42]. From this species, Kwok et al. [43] isolated the toxin trans-2-decenedioic acid, which is toxic to nematodes. Here, *P. chrysosporium* proved to be just as lethal to root-knot nematodes as *P. ostreatus*. We isolated *P. chrysosporium* strain B-22 as a nematophagous fungus lethal to the Southern root-knot nematode *M. incognita*. Previous studies on *P. chrysosporium* have focused on removing toxic environmental pollutants [44]. P*. chrysosporium* can produce lignin peroxidase, manganese peroxidase, and laccase [45], among others. Inoculation with *P. chrysosporium* can greatly reduce the disease index of wilt and root-knot nematode; thus, it might be promising to use it to overcome continuous cropping limitations, e.g., in cucumber[29]. Isolated *P. chrysosporium* has been used in the biodegradation of lignin and nicotine in tobacco stalk [46], as well as polycyclic aromatic hydrocarbons [45] and other pollutants [47,48]. Meanwhile, Xu et al. [29] used *P. chrysosporium* for the control of damping-off root wilt disease caused by the continuous cropping of cucumber. The present study identified *P. chrysosporium* as a parasitic fungus *infecting M. incognita*. The assay of the lethal effect by *P. chrysosporium* strain B-22 on the *M. incognita* J2 demonstrated that strain B-22 can control these nematodes. This assay provides the first evidence of parasitism of *P. chrysosporium* on *M. incognita*. To further confirm this case of parasitism, our study demonstrated that strain B-22 was parasitic on *M. incognita* at different life stages. This provided further strong evidence of the parasitism of *P. chrysosporium* on *M. incognita*.

In this study, *P. chrysosporium* strain B-22 parasitized the eggs of *M. incognita* by producing hyphae, which first surrounded the egg shell and then destroyed the eggs. Some studies have shown that the egg-parasitic fungi *Verticillium suchlasporium*, *P. lilacinus*, *Pochonia* spp., and *T. harzianum* can secrete protease and chitinase, which degrade certain cyst nematode proteins to effectively destroy the nematode egg shell and later parasitize and kill the eggs [49]. We speculated that the infection of *M. incognita* eggs by strain B-22 might also be involved in egg shell decomposition through the production of protease and chitinase.

*P. chrysosporium* strain B-22 parasitized the J2 of *M. incognita* by producing sticky conidia that enveloped them. These J2 were completely destroyed after 5 days of incubation. In general, the fungal conidia first stuck to the nematode cuticle or made some structures to adhere to. Then, massive hyphae grew and crossed and invaded the nematode cuticle, causing the deformation and death of J2 by leakage. Nematode cuticle decomposition indicates that *P. chrysosporium* strain B-22 might be able to produce proteases that degrade the nematode cuticle, as reported for *Lecanicillium psalliotae*, which produces an extracellular protease to degrade the nematode body [50]. Parasitic *P. ostreatus*, which is another white-rot fungus that attacks nematodes, was reported to produce toxin droplets to attack nematode J2, thereby killing them in 2h [24,42,51]. It is likely that *P. chrysosporium* strain B-22 employed a similar mechanism, i.e., produced toxin droplets to attack and kill the nematode.

Strain B-22 produced mycelial masses on the surface of the adult females of *M. incognita*; thus, the females were killed by the parasitizing mycelia. Our results indicate that *P. chrysosporium* strain B-22 can parasitize *M. incognita* and thus function as a biocontrol agent.

Results from our greenhouse experiments clearly demonstrated that different concentrations of *P. chrysosporium* strain B-22 could control different life stages of *M. incognita* and root knot in tomato. Moreover, the control efficacy increased with the conidial concentration of the B-22 suspension. In contrast, *P. chrysosporium* strain B-22 significantly increased plant height, root length, and aboveground and root fresh mass of tomato plants inoculated with *M. incognita*. Thus, *P. chrysosporium* strain B-22 effectively controlled *M. incognita*, while significantly improving the growth of tomato plants. These results indicate that *P. chrysosporium* strain B-22 is safe for greenhouse-grown tomato seedlings.

In conclusion, this study characterized the parasitism of *P. chrysosporium* strain B-22 on the eggs, J2, and adult females of the *M. incognita*. We also evaluated this strain’s control efficacy and safety when applied in tomato plant culture in a greenhouse. The results demonstrated the potential of *P. chrysosporium* strain B-22 as a promising biocontrol agent for the control of *M. incognita*. However, additional studies are needed to identify and characterize the molecules responsible for the toxic effect of this parasitic fungal strain on the nematode and the mechanism associated with the parasitic infection of *M. incognita*. In addition, this strain’s control efficacy under field conditions and against other root-knot nematodes needs to be investigated.

## Acknowledgments

This work was supported by the Shanxi Provincial Science and Technology Planning Project (Grant Nos. 20120311019-3, 20133054), The Key Research and Development Program Projects in Shanxi Province (Grant Nos. 201803D221004-2), and the Basic Research Program of Shanxi Province (Grant Nos. 201801D221321).

## Author contributions

Jianming Wang conceived and supervised the study; Yumei Xu designed the experiments; Bin Du and Li Yan performed the experiments; Hailong Dong analyzed the data; and Bin Du wrote the manuscript.

## References

1. Abad P, Gouzy J, Aury JM, Castagnone-Sereno P, Danchin EG (2008) Genome sequence of the metazoan plant-parasitic nematode Meloidogyne incognita. Nat Biotechnol 26: 909–915.

2. Jones JT, Haegeman A, Danchin EG, Gaur HS, Helder J, et al. (2013) Top 10 plant-parasitic nematodes in molecular plant pathology. Molecular plant pathology 14: 946–961.

3. Ding X, Shields J, Allen R, Hussey RS (2000) Molecular cloning and characterisation of a venom allergen AG5-like cDNA from Meloidogyne incognita. International Journal for Parasitology 30: 77–81.

4. Aocheng C, Meixia G, Dongdong Y, Liangang M, Qiuxia W, et al. (2014) Evaluation of sulfuryl fluoride as a soil fumigant in China. Pest Management Science 70: 219–227.

5. Trudgill, Blok (2001) Apomictic, polyphagous root-knot nematodes: exceptionally successful and damaging biotrophic root pathogens. Annu Rev Phytopathologia Mediterranea: 39: 53–77.

6. Sasser, Eisenback, Carter, Triantaphyllou (1983) The international Meloidogyne project - its goals and accomplishments. Annu Rev Phytopathol 21: 271–288.

7. Huang WK, Sun JH, Cui JK, Wang GF, Kong LA, et al. (2014) Efficacy evaluation of fungus Syncephalastrum racemosum and nematicide avermectin against the root-knot nematode Meloidogyne incognita on cucumber. Plos One 9: e89717.

8. Abo-Korah MS (2017) Biological Control of Root-Knot Nematode, Meloidogyne javanica Infecting Ground Cherry, Using Two Nematophagous and Mychorrhizal Fungi. Egyptian Journal of Biological Pest Control 27(1), 2017, 111–115.

9. Marin MV, Santos LS, Gaion LA, Rabelo HO, Franco CA, et al. (2017) Selection of resistant rootstocks to Meloidogyne enterolobii and M. incognita for okra (Abelmoschus esculentus L. Moench). Chilean journal of agricultural research 77: 57–64.

10. Mantelin S, Bellafiore S, Kyndt T (2017) Meloidogyne graminicola: a major threat to rice agriculture. Mol Plant Pathol 18: 3–15.

11. Chen J, Abawi GS, Zuckerman BM (1999) Suppression of Meloidogyne hapla and Its Damage to Lettuce Grown in a Mineral Soil Amended with Chitin and Biocontrol Organisms. Journal of Nematology 31: 719–725.

12. Ismail AE (2014) Growing Jatropha Curcas and Jatropha Gossypiifolia as a Interculture with Sunflower for Control of Meloidogyne Javanica in Egypt. International Journal of Sustainable Agricultural Research 1: 39–44.

13. Verdejo-Lucas S, Ornat C, Sorribas FJ, Stchiegel A (2002) Species of Root-knot Nematodes and Fungal Egg Parasites Recovered from Vegetables in Almería and Barcelona, Spain. Journal of Nematology 34: 405–408.

14. Ashraf MS, Khan TA (2005) Effect of opportunistic fungi on the life cycle of the root-knot nematode (Meloidogyne javanica) on brinjal. Archives of Phytopathology and Plant Protection 38: 227–233.

15. Trifonova Z, Karadjova J, Georgieva T (2009) Fungal parasites of the root-knot nematodes Meloidogyne spp. in southern Bulgaria. Estonian Journal of Ecology 58: 47–52.

16. Siddiqui ZA, Akhtar MS (2009) Effects of antagonistic fungi, plant growth-promoting rhizobacteria, and arbuscular mycorrhizal fungi alone and in combination on the reproduction of Meloidogyne incognita and growth of tomato. Journal of General Plant Pathology 75: 144.

17. Goswami J, Pandey RK, Tewari JP, Goswami BK (2008) Management of root knot nematode on tomato through application of fungal antagonists, Acremonium strictum and Trichoderma harzianum. Journal of Environmental Science & Health Part B 43: 237–240.

18. Vianene NM, Abawi GS (2000) Hirsutella rhossiliensisand Verticillium chlamydosporium as Biocontrol Agents of the Root-knot Nematode Meloidogyne hapla on Lettuce. Journal of Nematology 32: 85.

19. Anastasiadis IA, Giannakou IO, Prophetou-Athanasiadou DA, Gowen SR (2008) The combined effect of the application of a biocontrol agent Paecilomyces lilacinus, with various practices for the control of root-knot nematodes. Crop Protection 27: 352–361.

20. Li J, Zou C, Xu J, Ji X, Niu X, et al. (2015) Molecular Mechanisms of Nematode-Nematophagous Microbe Interactions: Basis for Biological Control of Plant-Parasitic Nematodes. Annual Review of Phytopathology 53: 67–95.

21. Zeng G-M, Chen A-W, Chen G-Q, Hu X-J, Guan S, et al. (2012) Responses of Phanerochaete chrysosporium to Toxic Pollutants: Physiological Flux, Oxidative Stress, and Detoxification. Environmental Science & Technology 46: 7818–7825.

22. Zeng G-M, Huang D, Huang G, Hu T, Jiang X, et al. (2007) Composting of lead-contaminated solid waste with inocula of white-rot fungus. Bioresource Technology 98: 320–326.

23. Kersten P, Cullen D (2007) Extracellular oxidative systems of the lignin-degrading Basidiomycete Phanerochaete chrysosporium. Fungal Genet Biol 44: 77–87.

24. Barron GL, Thor RG (1987) Destruction of nematodes by species of Pleurotus. Canadian Journal of Botany 65: 774–778.

25. Satou T, Kaneko K, Li W, Koike K (2008) The toxin produced by pleurotus ostreatus reduces the head size of nematodes. Biological & pharmaceutical bulletin 31: 574–576.

26. Iijima N, Yoshino H, Ten LC, Ando A, Watanabe K, et al. (2002) Two genes encoding fruit body lectins of Pleurotus cornucopiae: sequence similarity with the lectin of a nematode-trapping fungus. Biosci Biotechnol Biochem 66: 2083–2089.

27. Ray A, Ayoubi-Canaan P, Hartson SD, Prade R, Mort AJ (2012) Phanerochaete chrysosporium produces a diverse array of extracellular enzymes when grown on sorghum. Applied Microbiology and Biotechnology 93: 2075–2089.

28. Li P, Chen J, Li Y, Zhang K, Wang H (2017) Possible mechanisms of control of Fusarium wilt of cut chrysanthemum by Phanerochaete chrysosporium in continuous cropping fields: A case study. Scientific reports 7: 15994.

29. Xu SX, Zhang SM, You XY, Jia XC, Wu K (2008) Degradation of soil phenolic acids by Phanerochaete chrysosporium under continuous cropping of cucumber. Ying Yong Sheng Tai Xue Bao 19: 2480–2484.

30. Eisenback JD (1985) Detailed morphology and anatomy of second-stage juveniles, males, and females of the genus Meloidogyne (root-knot nematodes). In: Sasser JN, Carter CC, editors. An advanced treatise on meloidogyne. vol. 1: Biology and control. Raleigh, NC: North Carolina State University Graphics. 95–112.

31. Hussey RS, Barker KR (1973) A comparison of methods of collecting inocula of Meloidogyne species, including a new technique. Plant Dis Report 57: 1025–1028.

32. Burdsall HH, Jr., Eslyn WE (1974) A new Phanerochochaete with a Chrysosporium imperfect state. Mycotaxon 1: 123–133.

33. Mittal N, Saxena G, Mukerji KG (1995) Integrated control of root-knot disease in three crop plants using chitin and Paecilomyces lilacinus. Crop Protection 14: 647–651.

34. Zhang SW, Liu J, Xu BL, Gu LJ, Xue YY (2013) [Parasitic and lethal effects of Trichoderma longibrachiatum on Heterodera avenae: microscopic observation and bioassay]. Ying Yong Sheng Tai Xue Bao 24: 2955–2960.

35. Schwartz HT (2007) A protocol describing pharynx counts and a review of other assays of apoptotic cell death in the nematode worm Caenorhabditis elegans. Nat Protoc 2: 705–714.

36. Dong H, Zhou X-G, Wang J, Xu Y, Lu P (2015) Myrothecium verrucaria strain X-16, a novel parasitic fungus to Meloidogyne hapla. Biological Control 83: 7–12.

37. Yan XN, Sikora RA, Zheng JW (2011) Potential use of cucumber (Cucumis sativus L.) endophytic fungi as seed treatment agents against root-knot nematode Meloidogyne incognita. J Zhejiang Univ Sci B 12: 219–225.

38. Castillo, Nico, Azcón-Aguilar (2006) Protection of olive planting stocks against parasitism of root-knot nematodes by arbuscular mycorrhizal fungi. Plant Pathology 55: 705–713.

39. Sharon E, Chet I, Viterbo A, Bar-Eyal M, Nagan H, et al. (2007) Parasitism of Trichoderma on Meloidogyne javanica and role of the gelatinous matrix. European Journal of Plant Pathology 118: 247–258.

40. Bird DM, Kaloshian I (2003) Are roots special? Nematodes have their say. Physiological and Molecular Plant Pathology 62: 115–123.

41. Affokpon A, Coyne DL, Htay CC, Agbèdè RD, Lawouin L, et al. (2011) Biocontrol potential of native Trichoderma isolates against root-knot nematodes in West African vegetable production systems. Soil Biology and Biochemistry 43: 600–608.

42. Thorn RG, Barron GL (1984) Carnivorous mushrooms. Science 224: 76–78.

43. Kwok OCH, Plattner, Weisleder, Wicklow D (1992) A nematicidal toxin fromPleurotus ostreatus NRRL 3526. Journal of Chemical Ecology 18: 127–137.

44. Tan Q, Chen G, Zeng G, Chen A, Guan S, et al. (2015) Physiological fluxes and antioxidative enzymes activities of immobilized Phanerochaete chrysosporium loaded with TiO2 nanoparticles after exposure to toxic pollutants in solution. Chemosphere 128: 21–27.

45. Kadri T, Rouissi T, Kaur Brar S, Cledon M, Sarma S, et al. (2017) Biodegradation of polycyclic aromatic hydrocarbons (PAHs) by fungal enzymes: A review. J Environ Sci (China) 51: 52–74.

46. Su Y, Xian H, Shi S, Zhang C, Manik SM, et al. (2016) Biodegradation of lignin and nicotine with white rot fungi for the delignification and detoxification of tobacco stalk. BMC Biotechnol 16: 81.

47. Hu L, Zeng G, Chen G, Dong H, Liu Y, et al. (2016) Treatment of landfill leachate using immobilized Phanerochaete chrysosporium loaded with nitrogen-doped TiO(2) nanoparticles. J Hazard Mater 301: 106–118.

48. Herve V, Ketter E, Pierrat JC, Gelhaye E, Frey-Klett P (2016) Impact of Phanerochaete chrysosporium on the Functional Diversity of Bacterial Communities Associated with Decaying Wood. PLoS One 11: e0147100.

49. Wei B-Q, Xue Q-Y, Wei L-H, Niu D-D, Liu H-X, et al. (2009) A novel screening strategy to identify biocontrol fungi using protease production or chitinase activity against Meloidogyne root-knot nematodes. Biocontrol Science and Technology 19: 859–870.

50. Yang J, Huang X, Tian B, Wang M, Niu Q, et al. (2005) Isolation and characterization of a serine protease from the nematophagous fungus, Lecanicillium psalliotae, displaying nematicidal activity. Biotechnol Lett 27: 1123–1128.

51. Thorn G, Tsuneda A (1993) Interactions between Pleurotus species, nematodes, and bacteria on agar and in wood. Transactions of the Mycological Society of Japan 34: 449–464.

